# Homozygous *STAT2* gain-of-function mutation by loss of USP18 activity in a patient with type I interferonopathy

**DOI:** 10.1101/2019.12.12.874123

**Authors:** Conor Gruber, Marta Martin-Fernandez, Fatima Ailal, Xueer Qiu, Justin Taft, Jennie Altman, Jéremie Rosain, Sofija Buta, Aziz Bousfiha, Jean-Laurent Casanova, Jacinta Bustamante, Dusan Bogunovic

## Abstract

Type I interferonopathies are monogenic disorders characterized by enhanced Type I interferon (IFN-I) activity. Inherited ISG15 and USP18 deficiencies underlie type I interferonopathies by preventing the regulation of late responses to IFN-I. Specifically, ISG15/USP18 are induced by IFN-I and sterically hinder JAK1 from binding to the IFNAR2 subunit of IFN-I receptor. We report an infant who died of autoinflammation due to a homozygous missense mutation (R148Q) in *STAT2*. The variant is gain-of-function (GOF) for ISGF3-dependent induction of late but not early response to IFN-I. Surprisingly, the mutation does not enhance the intrinsic transcriptional activity of ISGF3. Rather, the STAT2 R148Q variant is GOF because it fails to appropriately interact with and traffic USP18 to IFNAR2, preventing USP18 from negatively regulating responses to IFN-I. Overall, a STAT2 missense mutation that fails to facilitate USP18-mediated signal termination in the homozygous state underlies a novel genetic etiology of type I interferonopathy.

## Introduction

Human type I interferons (IFN-is) form a group of 19 potent antiviral and proinflammatory cytokines (Borden et al., 2007). They are produced by almost any cell type in response to various stimuli, particularly viral intermediates and by-products (Kawai and Akira, 2008), although some IFN-I subtypes seem to be preferentially produced by certain cell types (Marks et al., 2019). Once produced, IFN-Is are secreted, and, in an autocrine and paracrine fashion, signal through the IFN-I receptor (IFNAR), which consists of two subunits, IFN-I receptor I (IFNAR1) and IFN-I receptor II (IFNAR2). Most, if not all, cells express the IFN receptor. Upon ligand binding, the IFN-I-IFNAR complex then initiates a signaling cascade via auto- and transphosphorylation of JAK1 and TYK2, kinases that subsequently phosphorylate STAT1 and STAT2. In complex with IRF9, STAT1/2 form ISGF3 that translocates to the nucleus and initiates transcription of hundreds of IFN-I stimulated genes (ISGs) (Schindler et al., 2007; Stark and Darnell, 2012). While IFN-I cytokines don’t directly activate transcription of IFN-I genes, they potentiate their own production by increasing the sensors or transcription factors capable of inducing IFN-I genes, in particular via IRF7 (Levy et al., 2002). In contrast, two ISGs, ISG15 (Zhang et al., 2015) and USP18 (Meuwissen et al., 2016), form a complex which negatively regulates response to IFN-I. Namely USP18, which when bound by ISG15 is more stable, displaces JAK1 (Francois-Newton et al., 2012) from IFNAR2, thereby terminating responses to and amplification of IFN-I.

Dysregulation of IFN-I activity has dire consequences for humans. Patients with autosomal recessive, complete *IFNAR1* (Hernandez et al., 2019), *IFNAR2* (Duncan et al., 2015), *JAK1* (Eletto et al., 2016), *TYK2* (Kreins et al., 2015), *STAT1* (Dupuis et al., 2003), *STAT2* (Hambleton et al., 2013), and *IRF9* (Hernandez et al., 2018) deficiencies suffer from severe infectious diseases. Patients with IFNAR1, IFNAR2, STAT2, and IRF9 deficiencies are exclusively prone to viral diseases (Duncan et al., 2015; Hambleton et al., 2013; Hernandez et al., 2019; Hernandez et al., 2018), while patients with TYK2 and STAT1 deficiencies are also prone to mycobacterial disease (Del Bel et al., 2017; Kreins et al., 2015; Wu and Holland, 2015). In contrast, Type I Interferonopathies are monogenic disorders that result from autoinflammation caused by excessive IFN-I activity (Rodero and Crow, 2016). The concept was established, and the term was coined by, Yanick Crow (Crow, 2011; Rodero and Crow, 2016). Known etiologies disrupt either of two distinct cellular mechanisms. The *production* of IFN-I can be excessive owing to bi-allelic loss-of-function (LOF) mutations in self-RNA and DNA digesting enzymes or mono-allelic gain-of-function (GOF) mutations in RNA and DNA cytoplasmic sensors (Rodero and Crow, 2016). Alternatively, there can be hyperactive *responses* to IFN-I by bi-allelic LOF mutations in negative regulators of IFN-I, such as ISG15 (Zhang et al., 2015) and USP18 (Meuwissen et al., 2016), or by heterozygous GOF mutations in JAK1 (Del Bel et al., 2017; Gruber et al., 2019) and STAT1 (Liu et al., 2011). Herein, we studied a child who died of unexplained, severe, early-onset type I interferonopathy.

## Results

### *Ex vivo* phenotyping reveals type I interferonopathy

We studied a patient (P1, II.9) born to consanguineous patients from Morocco who presented with an early-onset, severe autoinflammation evocative of severe type I interferonopathy. He presented with skin ulcerations, seizures, cerebral calcifications, and ultimately respiratory failure and death (please see material and methods for clinical details, Figure 1A). We first measured whole blood mRNA levels of four *ISGs*, including *IFIT1, IFI27, RSAD2* and *ISG15*, all of which were elevated nearly 1000-fold as compared to the healthy donor or the heterozygous mother (Figure 1B). Furthermore, using SiMoA, a digital ELISA that allows for attomolar sensitivity in IFN-α detection, we detected drastically high levels of IFN-α in patient plasma (1,000 fg/ml as compared to an average 50 fg/ml in healthy controls) (Figure 1C). Given these classical features of type I interferonopathy in immune peripheral cells (Rodero and Crow, 2016), we then examined the composition of peripheral blood mononuclear cells (PBMCs). We detected an aberrant distribution of immune cell subtypes, with proportionally less myeloid and NK cells and more B and T cells, suggesting a severely dysregulated immunophenotype (Figure 1D-E, Supplementary Table I). Given the overt IFN-I signature, we took advantage of surface expression of SIGLEC1 (CD169), an ISG, and determined that myeloid cells, in particular classical monocytes (CD14^+^CD16^low^), as well as dendritic cells, had significantly augmented expression of CD169 (Figure 1F). This perhaps suggested that the most affected were myeloid cells, and that they largely contribute to the ISG signature commonly monitored in whole blood. Combined, this *ex vivo* analyses confirmed the *bona fide* type I interferonopathy.

**Figure 1.**
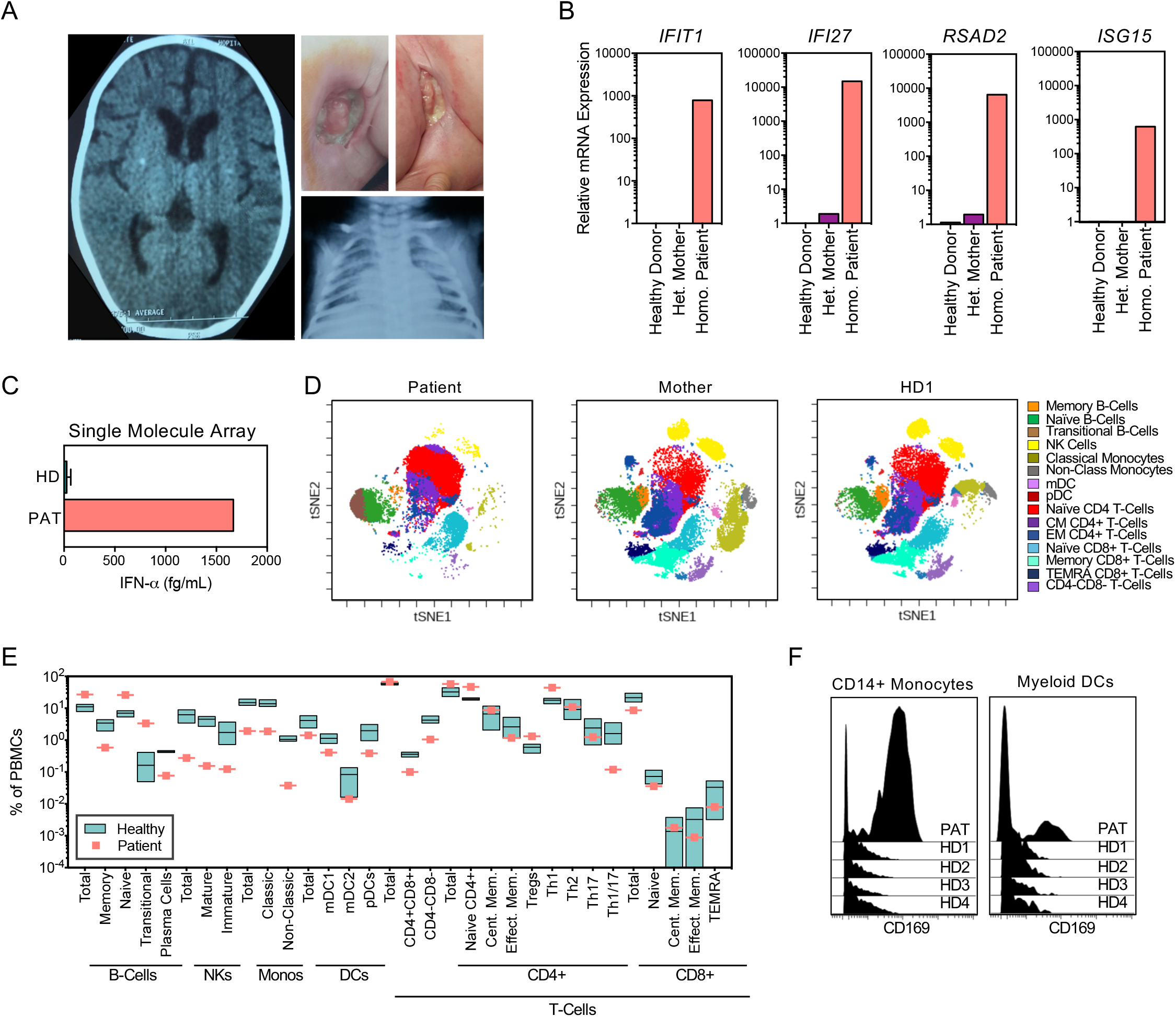
Peripheral blood signature indicates autosomal recessive type I interferonopathy. (A) Clinical signs of immune pathology, clockwise from top: CT scan demonstrating calcifications of the frontal and parietal lobes, fistulizing adenitis of the axillary and inguinal nodes, chest radiograph displaying bilateral opacities of the lungs. (B) mRNA expression of interferon stimulated genes measured from whole blood RNA isolated from healthy donor, the patient’s mother and the patient. (C) Quantification of circulating IFN-alpha by digital ELISA (Single Molecule Array) in plasma from three healthy controls (HD) and the patient (PAT). (D) tSNE plots demonstrating the immunophenotyped of peripheral blood mononuclear cells (PBMCs) as determined by mass cytometry. (E) Quantification of immune cells populations of 4 healthy donors and the patient expressed as percent of total PBMCs. (F) Histograms of CD169 expression, an interferon stimulated gene, in classical monocytes and myeloid DCs.

### Whole exome sequencing reveals a homozygous mutation in *STAT2*

We performed whole-exome sequencing (WES) of P1 and searched for candidate genetic variants, testing an autosomal recessive (AR) model, given the consanguinity and history of prior infant deaths in the family (Figure 2A). A homozygous variant was identified in exon 5 of the signal transducer and activator of transcription 2 (STAT2) at position c.443G>A, which results in the substitution of the arginine at position 148 by a glutamine, p.R148Q (Figure 2A-B). No other putative disease-causing alleles were detected in any of 19 known genes responsible of interferonopathies (Rodero and Crow, 2016)(Supplementary Figure 1A). There were eight other homozygous rare non-synonymous variations, none of which was in a gene known to be related to IFN-I (Supplemental Table 3). Sanger sequencing of exon 5 of *STAT2* confirmed the homozygous mutation of genomic DNA from whole blood and fibroblasts from P1 (Figure 2C). DNA from the mother was heterozygous for this variant, while the father’s DNA was unavailable. *In silico* analysis revealed that the variant was predicted to be protein-damaging by combined annotation-dependent depletion (CADD,score = 24.1), above the mutation significance cutoff (MSC) of 2.313 (Supplemental Figure 1B). The variant is located in the coiled-coil domain of STAT2 at a residue (148) that is highly conserved across mammals, but not in rodents (Figure 2D). This variant is absent in our in-house database of over 6,000 WES and in public databases (the GME variome or in the gnomAD v2.1). There are only eight non-synonymous *STAT2* variations found in homozygosity in public databases, none of which is predicted to be LOF. The homozygous mutations in *STAT2* that were previously shown to be loss-of-expression and LOF in terms of ISGF3 activity in patients with severe viral disease were private (Hambleton et al., 2013; Moens et al., 2017). Altogether, these findings suggested that the c.443G>A allele may be disease-causing. As patients homozygous for loss-of-expression STAT2 mutations had impaired responses to IFN-I and viral diseases, our findings further suggested that P1 might be homozygous for a *STAT2* mutation that is GOF for ISGF3 activity, or LOF for a function other than ISGF3 activity, or both.

**Figure 2.**
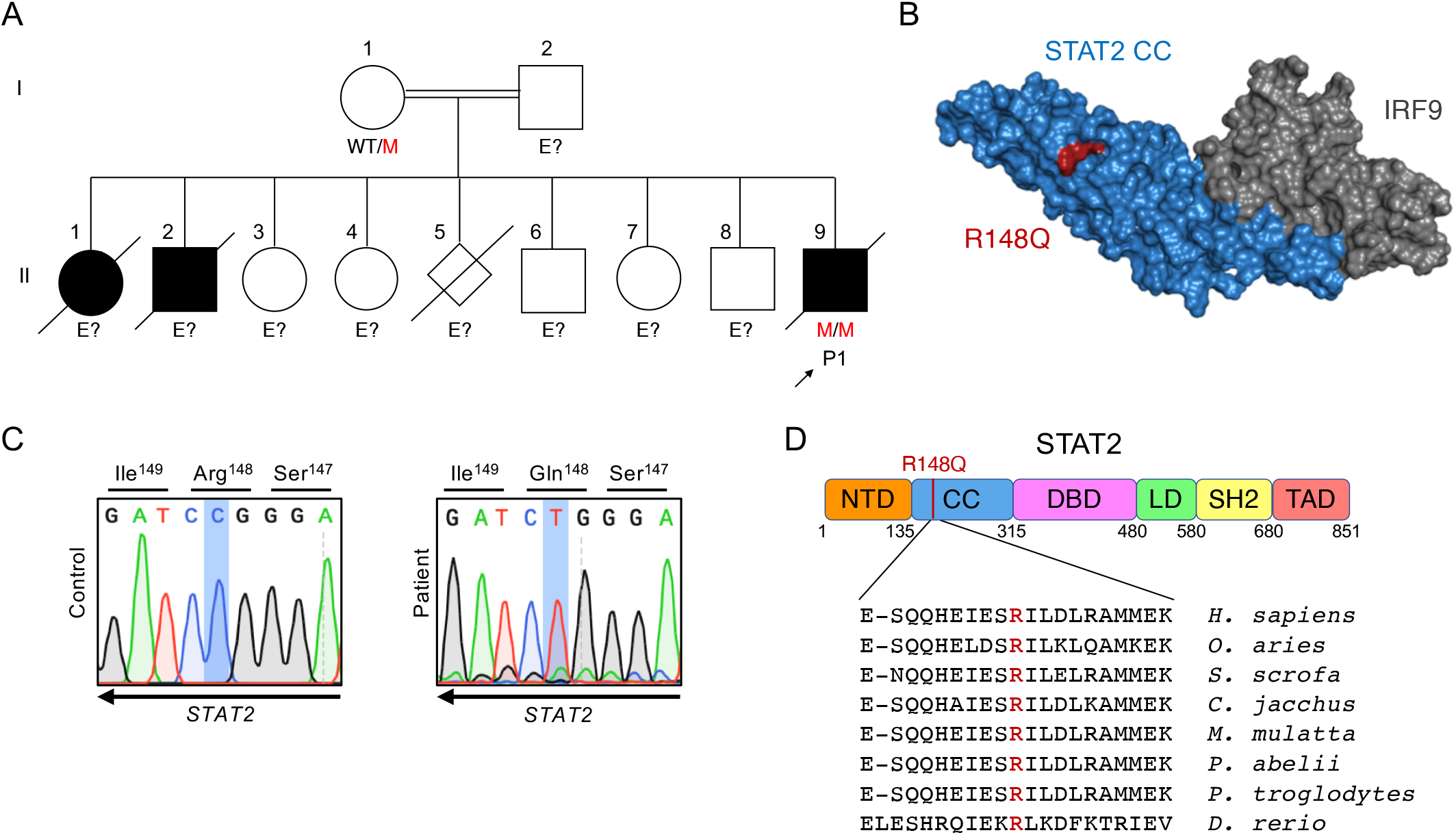
Homozygous mutation affecting the coiled-coil domain of STAT2 identified in a patient with lethal autoinflammation. (A) Family pedigree displaying affected individuals (shaded) and genotypes at position c.443 in *STAT2*. (B) Crystal structure of murine STAT2 (blue) in complex with IRF9 (gray), demonstrating position of R148Q (red) in the coiled-coil domain (CC), but outside of the STAT2/IRF9 interface. (C) Sanger sequencing of genomic DNA from patient and healthy donor fibroblasts. (D) Schematic of STAT2 structure, indicating localization of mutated residue and its conservation across mammals.

### The R148Q STAT2 variant is GOF for ISGF3 activity

In order to characterize the allele in isolation of the patient’s genetic background, we took advantage of U6A cells, a fibrosarcoma human cell line that is STAT2 null. We transduced U6A cells with either *Luciferase* (negative control), wild type (WT) *STAT2* or R148Q *STAT2*. We first detected that transduction with either WT or R148Q *STAT2* resulted in similar levels of *STAT2* mRNA and STAT2 protein (Figure 3A and B). In addition, brief 4-hour stimulation with IFN-I resulted in similar mRNA levels of *MX1* and *IFI27* (Supplementary Figure 1C). In contrast, prolonged 16-hour stimulation with IFN-I, resulted in increased mRNA levels of *MX1, IFI27* and *IFIT1* in U6A cells transduced with R148Q STAT2, when compared with WT STAT2 (Figure 3B). Prolonged stimulation with Type II IFN, which does not utilize STAT2, was normal (Figure Supplement 1D). We then hTert-immortalized and tested dermal fibroblasts derived from the patient. Expression of *MX1*, *RSAD2* and *IFIT1* mRNA was significantly elevated in patient hTert fibroblasts stimulated with IFNα2b when compared with cells from healthy controls at late time points (Figure 3D). We tested whether this phenotype requires homozygosity. We co-expressed WT and R148Q STAT2 in U6A cells. In this system, upon IFN-I stimulation, the presence of WT STAT2 rescued the heightened signaling afforded by R148Q STAT2 (Figure 3D), whereas transduction with the negative control did not. Cells from heterozygous relatives of the patient were not available. They were, however, healthy and had no elevation of ISGs in peripheral blood (Figure 1B), suggesting that the GOF mutation likely has no impact in heterozygosity. Collectively, these experiments indicate that the *STAT2* mutation is GOF in isolation and that the bi-allelic genotype also results in gain of ISGF3 activity.

**Figure 3.**
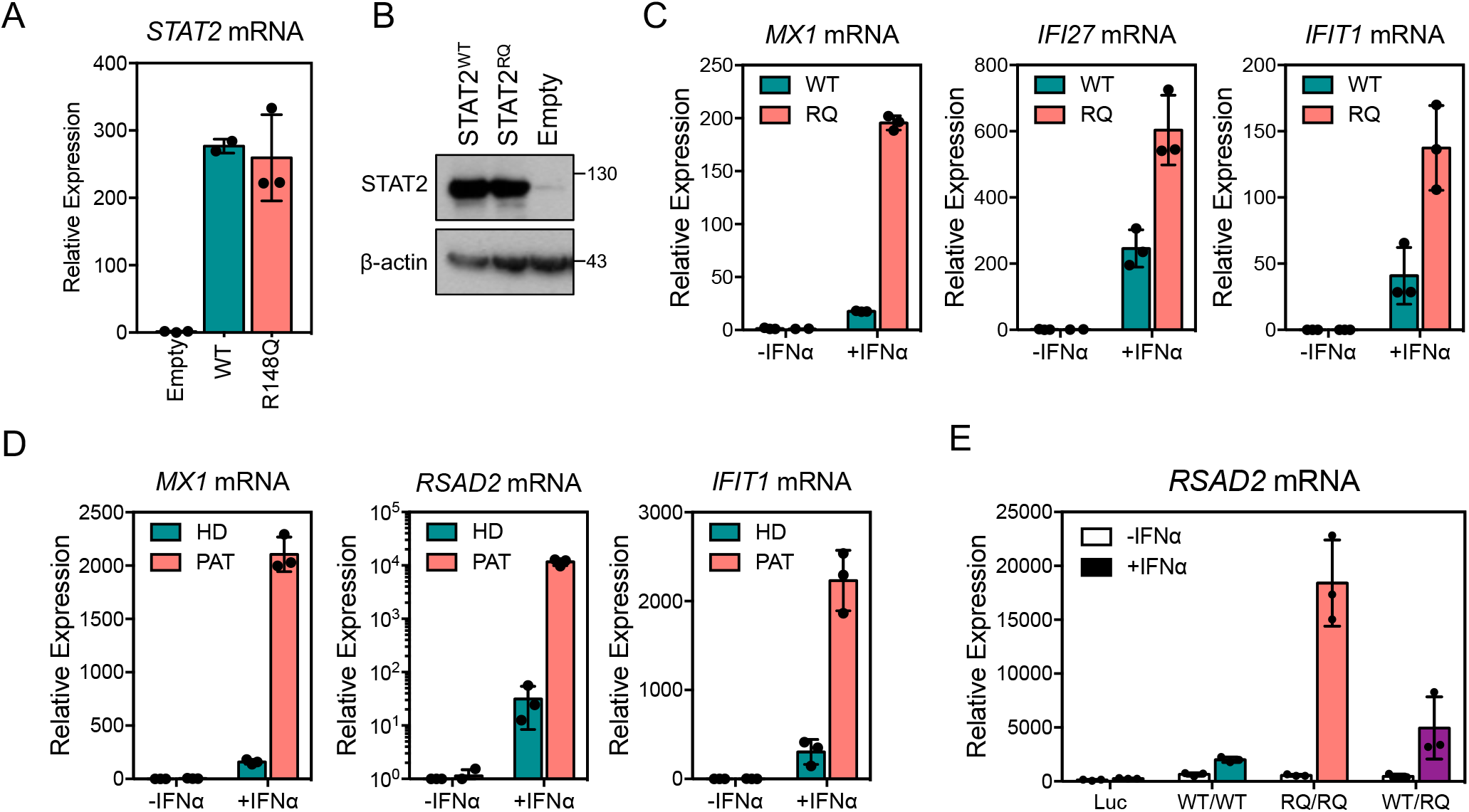
STAT2 R148Q leads to hyper-induction of interferon-stimulated genes (ISGs), which can be rescued by WT STAT2. (A) *STAT2* transcript levels in U6A cells (STAT2−/−) reconstituted with an empty vector, WT *STAT2* (WT) or R148Q *STAT2* (RQ). (B) STAT2 protein levels by western blot in HEK239T cells transfected with indicated plasmids. (C) Expression of interferon stimulated genes (ISGs) in transduced U6A cells after 16 hours of 100 IU/mL interferon-α stimulation. (D) Induction of ISGs in hTERT-immortalized fibroblasts. (E) Transfection of WT *STAT2* (WT), R148Q *STAT2* (RQ) or both in U6A cells, followed by overnight stimulation with interferon-α. All results are representative of at least 3 independent experiments.

### LOF of STAT2-mediated regulation of USP18 results in overall GOF phenotype in R148Q STAT2

Two functions of STAT2 are known. As part of the ISGF3 transcription factor complex, STAT2 is essential for cellular responses to IFN-I and IFN-III downstream from their respective receptors. STAT2 was also recently discovered to shuttle USP18 to IFNAR2, where USP18 exerts its negative regulatory function of IFN-I responses (Arimoto et al., 2017). STAT2 is therefore a positive regulator of IFN-I, while it also contributes to the negative regulation of IFN-I once USP18 is produced. Given that R148Q is located within the STAT2/USP18 interacting domain (Arimoto et al., 2017), we hypothesized that it retains positive ISGF3 activity but loses negative USP18-related regulatory function. We first tested proximal signaling in healthy control and patient hTert fibroblasts by stimulating cells with 30 min 1000 IU IFNα2b pulse. Both STAT1 and STAT2 phosphorylation were normal (Figure 4A), which was also the case in U6A cells (Supplemental Figure 1E-F). Nuclear localization after stimulation (Figure 4B) and subsequent nuclear export (Supplemental Figure 1G) were also normal for the mutant STAT2. We then tested STAT2 dephosphorylation in the immediate hours after IFN-I exposure, which appeared equal between WT and patient cells (Figure 4C). We also determined the ability of WT and R148Q STAT2 to immunoprecipitate with USP18. The mutant protein exhibited similar affinity towards USP18, indicating that although interaction was still possible, function was likely perturbed (Figure 4D). We then evaluated USP18 localization to IFNAR2, in presence of WT and R148Q STAT2. By immunoprecipitation of the receptor complex, we detected a reduced ability of R148Q STAT2 to recruit USP18 to IFNAR2 (Figure 4E). To functionally access this improper STAT2-USP18 homing, we tested the negative regulatory function of USP18/STAT2 R148Q by its capacity to prevent continual signaling. In cells WT for STAT2, stimulation with IFN-I leads to induction of USP18, which then significantly attenuates secondary challenge with IFN-I. While this was indeed the case in WT hTert fibroblasts, patient fibroblasts were unable to attenuate proximal signaling upon IFN-I restimulation, despite accumulating augmented levels of USP18 (Figure 4F). This effect was also observed in the U6A system (Supplemental Figure 1H). Collectively these results suggest a defect in USP18 trafficking to the receptor and explains both the GOF and recessive inheritance of *STAT2* R148Q.

**Figure 4.**
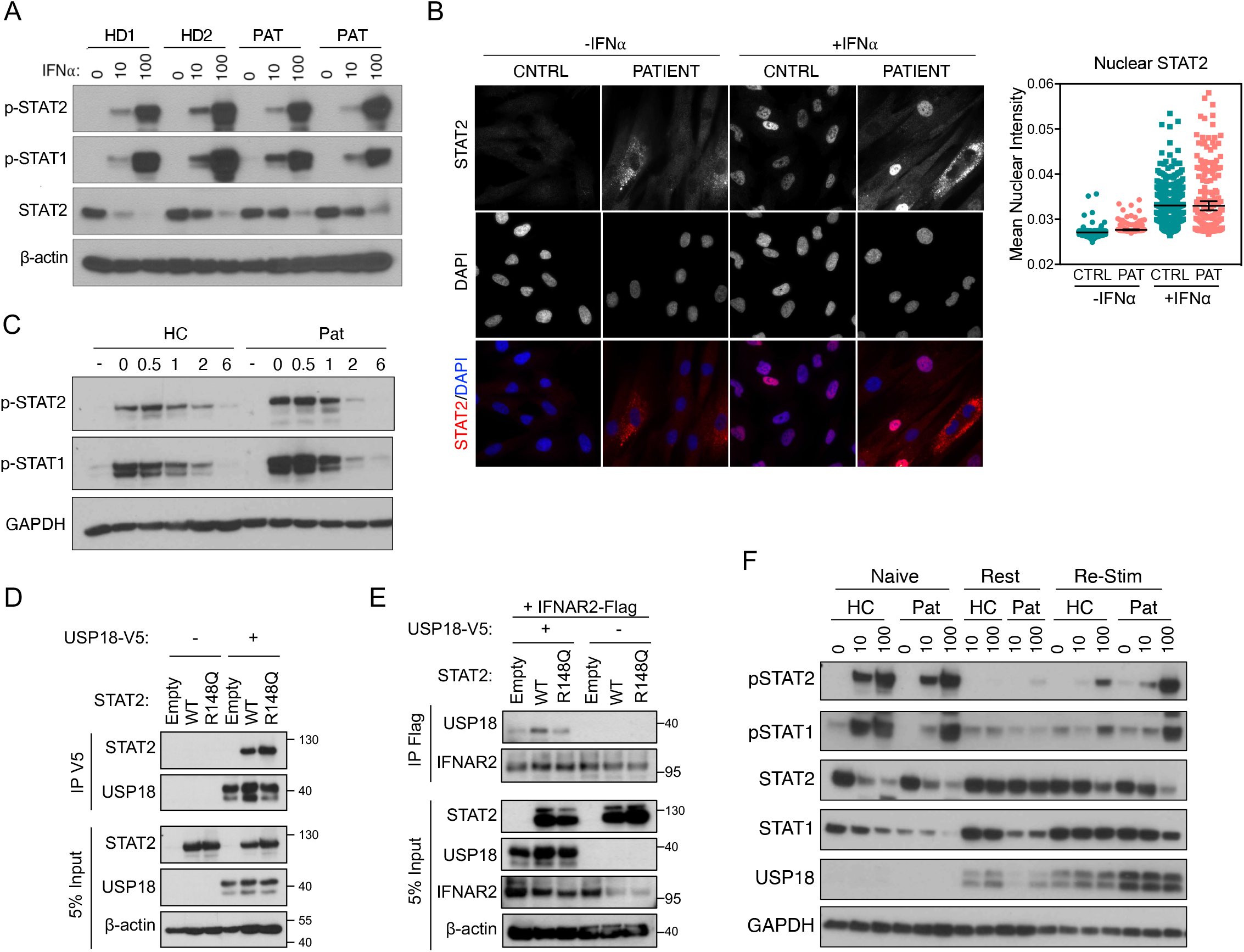
Intact proximal but not late signaling with STAT2 R148Q owing to defective USP18-mediated negative regulation. (A) Analysis of STAT1 and STAT2 phosphorylation in dermal fibroblasts following 15-minute stimulation with indicated doses (IU/mL) of IFN-α. Separate healthy donors (HD1, HD2) and two biological replicates of patient fibroblasts were tested. (B) Immunofluorescence staining of STAT2 translocation in fibroblasts after 30-minute stimulation with 1000 IU/mL IFN-α. Nuclear translocation quantified in right panel. (C) De-phosphorylation of STAT1 and STAT2 in hTERT fibroblasts pulsed with IFN-α for 15-minutes, washed and allowed to rest. (D) Transfection of indicated plasmids into HEK293T cells and subsequent immunoprecipitation of USP18-V5 to measure interaction with STAT2. (E) Recruitment of USP18 to the receptor in U6A cells transduced with either negative control, WT STAT2 or mutant STAT2, and transfected with IFNAR2-Flag and USP18-V5. Immunoprecipitation performed against IFNAR2-Flag. (F) (G) Analysis of negative regulatory capability by stimulating cells which had previously been stimulated (re-stim) or not (naive) with a primary stimulus of IFN-alpha for 12 hours, wash, and allowed to rest for 6 hours (rest) or 36 hours before a re-stimulation with indicated doses of IFN-α. All results are representative of 2-3 independent experiments.

## Discussion

Historically, the study of inborn errors of human immunity has provided invaluable insight into the intricate regulation of the immune system. To date, documented homozygous mutations affecting the IFN-I signaling cascade have exclusively been LOF and hypomorphic, while heterozygous mutations resulted in either dominant negative, haploinsufficient or GOF outcomes. In IFN-I signaling cascade, a homozygous GOF variant has never been reported until this discovery. To our knowledge, there are only two other documented homozygous GOF mutations described in genetics: in hypocalcemic hypoparathyroidism caused by homozygous mutations in *CASR* (Cavaco et al., 2018), and most recently in recurrent respiratory papillomatosis caused by homozygous mutations in *NLRP1* (Drutman et al., 2019). Identification of homozygous GOF mutation in *STAT2* was surprising, given that STAT2 is a positive regulator of the IFN-I pathway by its essential role in the transcription factor complex that induces hundreds of ISGs. However, STAT2 has a secondary role. Via its DNA binding and coiled-coil domains it binds and recruits USP18, an IFN-I negative regulator, to IFNAR2 (Arimoto et al., 2017). Thus, for a homozygous GOF mutations in STAT2 to exist, mutations would affect USP18 but not ISGF3 activity, as documented here with R148Q STAT2. For these reasons, homozygous GOF mutations in *STAT2* will likely remain exquisitely rare, as the pathogenic mutations must perturb the negatively regulatory domains while sparing the principal functions.

The patient reported here suffered from severe early onset inflammation characteristic of Type I Interferonopathies. STAT2 deficiency, alongside ISG15 and USP18 deficiencies, now constitutes a third genetic etiology leading to inadequate control of response to IFN-I. Of note, this novel disease largely phenocopies USP18 deficiency in clinical presentation and molecular mechanism (Alsohime et al., *In Press*; Meuwissen et al., 2016). Clearly, tight control of IFN-I receptor is essential for viability. If diagnosed and treated rapidly, as was the case in a recent USP18-deficient child, perhaps JAK inhibitor therapy would have rescued the life of this unfortunate child and deceased siblings (Alsohime et al., *In Press*). This possibility is further substantiated by several recent studies that successfully introduced JAK inhibitors in a diverse range of genetic etiologies of type I Interferonopathies (Sanchez et al., 2018). In the future, genetic diagnosis of rare *STAT2* mutations, especially in the USP18 interacting domain, should alert clinicians towards fast therapeutic interventions with current and new classes of JAK inhibitors, as they can indeed be lifesaving.

## Acknowledgements

We thank Adeeb Rahman, Brian Lee, Kevin Tuballes from Human Immune Monitoring Center at the Icahn School of Medicine for their technical assistance and guidance in experimental design. We also thank Ludovic Debure and Thomas Wisniewski for their technical assistance on the SiMoA ELISA assay. This research was supported by National Institute of Allergy and Infectious Diseases Grants R01AI127372, R21 AI134366 and R21AI129827, and funding from the March of Dimes, awarded to DB. CG was supported by T32 training grant 5T32HD075735-07. The Laboratory of Human Genetics of Infectious Diseases is supported by grants from the St. Giles Foundation, the Jeffrey Modell Foundation, The Rockefeller University Center for Clinical and Translational Science grant number 8UL1TR000043 from the National Center for Research Resources and the National Center for Advancing Sciences (NCATS), National Institutes of Health, the National Institute of Allergy and Infectious Diseases (5R01AI089970-02 and 5R37AI095983), grants from the Integrative Biology of Emerging Infectious Diseases Laboratory of Excellence (ANR-10-LABX-62-IBEID), Foundation for Medical Research (FRM, Fondation pour la Recherche Médicale) and the French National Research Agency (ANR) under the “Investments for the future” program (ANR-10-IAHU-01), GENMSMD (ANR-16-CE17-0005-01), *Institut National de la Santé et de la Recherche Médicale* (INSERM), Paris Descartes University and The Rockefeller University.

## Author contributions

CG, MMF designed and performed most of the experiments, analyzed the data and wrote the manuscript. JT helped design and perform microscopy. XQ performed IP experiments. SF helped with cloning, tissue culture and experiment execution. FA provided samples and performed clinical diagnoses and follow-up of the kindred. AB, JB and JLC provided patient samples, analyzed, and/or interpreted data, and contributed to the writing of the paper. DB helped design the experiments and analyze data, supervised the work and wrote the manuscript. All authors commented on the manuscript and approved its final version.

## Methods

### Genomic DNA extraction and Whole Exome Sequencing (WES)

Genomic DNA was isolated from the whole blood of patient, mother and healthy donors with the iPrep instruments from Thermo Fisher Scientific. Three micrograms of DNA were used for generation of WES from P1. The Agilent 50 Mb SureSelect exome kit was used in accordance with the manufacturer’s instructions. BWA aligner was used to align the reads with the human reference genome hg19, before recalibration and annotation with GATK, PICARD and ANNOVAR. Filters of variants was achieved with our in-house software. Mutation was verified by Sanger methods (PCR amplification conditions are available upon request). PCR products were analyzed by electrophoresis in 1% agarose gels, sequenced with V3.1 Big Dye terminator cycle sequencing kit and analyzed on an ABI Prism 3700 machine (Applied Biosystems, Foster City, CA).

### Cell Culture

HEK293T, U6A and hTert-immortalized dermal fibroblasts from the patient were cultured in DMEM (Gibco) supplemented with 10% fetal bovine serum (FBS) (Invitrogen), GlutaMAX (350 ng/ml; Gibco), and penicillin/streptomycin (Gibco). All cells were cultured at 37°C and 10% CO_2_. All cell lines were tested for mycoplasma contamination with the MycoAlert(tm) PLUS Mycoplasma Detection Kit (Lonza) according to the manufacturer’s instructions. U6A cells (STAT2−/−) were gifted by Dr. Sandra Pellegrini, Institut Pasteur.

### RNA Isolation and qPCR

All cytokine stimulations were performed as indicated with Interferon-alpha 2b (Intron-A) or Interferon-gamma (Biolegend, 570202) in complete DMEM. RNA was extracted from U6A cells, hTERT-immortalized fibroblasts (Qiagen RNeasy) or whole blood (PAXgene Blood RNA Kit) and reverse-transcribed (ABI High Capacity Reverse Transcriptase). The expression of ISGs (*IFIT1, MX1*, *RSAD2, IFI27, ISG15*), relative to the *18S* housekeeper gene, was analyzed by Taqman quantitative real-time PCR (TaqMan Universal Master Mix II w/ UNG) on a Roche LightCycler 480 II. The relative levels of ISG expression were calculated by the ΔΔCT method, relative to the mean values for the mock-treated controls or healthy donor.

### Ectopic Expression

HEK293T cells were transiently transfected with Lipofectamine 2000 (Thermoscientific) complexed with different constructs according to manufacturer’s instructions. The following genes-constructs were used for expression and co-immunoprecipitation: pTRIP-*USP18*-V5, PLX304-*IFNAR2*, pTRIP-*STAT2* (WT and R148Q). R148Q mutation was generated by site-directed mutagenesis on a WT *STAT2* plasmid using QuickChange II PCR (Agilent). Lentiviral particles were generated by co-transfection of *STAT2*, ps*PAX2* and p*MD2* by CaCl_2_ transfection. Supernatants were collected 48 hours later, purified and transferred to target cells with polybrene. Cells were selected with puromycin (0.4 micrograms/mL). To compensate for equivalent levels of ectopic expression, transfection and transduction experiments were matched with an irrelevant gene (*Luciferase*) in the same plasmid construct.

### Protein assays

Whole-cell extracts for immunoblotting were prepared by incubating cells for 10 min in RIPA lysis buffer (Thermo Fisher Scientific) with 50mM DTT and Protease/Phophatase inhibitor cocktail (Cell Signaling Technology). For co-immunoprecipitation assay, cells were lysed in 50 mM Tris pH 6.8, 0.5 % NP-40, 200 mM NaCl, 10% glycerol, 1 mM EDTA and 1x Protease/Phophatase inhibitor cocktail (Cell Signaling Technology). Cell lysates were incubated with V5 conjugated with protein G dynabeads for 2 hr at room temperature. Immunoprecipitates were subjected to western blotting. Immunoblotting was performed using the BioRad western blot workflow. Membranes were blocked in 5% BSA for primary antibodies or 5% nonfat dry milk for secondary antibodies. Antibodies used: STAT1 (Santa Cruz Biotechnology), STAT2 (Millipore), phospho-Tyr 701 STAT1 (Cell Signaling Technology), phospho-Tyr 689 STAT2 (Millipore), USP18 (Cell Signaling Technology), β-actin (ABclonal), and GAPDH (Millipore). Signal was detected with enhanced chemiluminescence detection reagent (SuperSignal West pico, Thermoscientific) by film development.

### Mass Cytometry

Whole blood from the patient and healthy controls were subject to Ficoll gradient to collect the mononuclear cell layer (PBMC). PBMCs were stained and analyzed by mass cytometry (CyTOF) at the Human Immune Monitoring Center of the Icahn School of Medicine at Mt. Sinai. Samples were barcoded then stained together with antibodies against selected surface markers for 30 minutes on ice. Cells were then washed and fixed, resuspended in diH_2_O containing EQ Four Element Calibration Beads (Fluidigm) and acquired on a CyTOF2 Mass Cytometer (Fluidigm). Data files were normalized by using a bead-based normalization algorithm (CyTOF software, Fluidigm) and debarcoded using CD45 gating. The gated populations were visualized in lower dimensions using viSNE in Cytobank (https://www.cytobank.org/) and manually gated based on the following traditionally-defined markers: CD45-Y89, CD57-In113, CD11c0In115, CD33-Pr141, CD19-Nd142, CD45RA-Nd143, CD141-Nd144, CD4-Nd145, CD8-Nd146, CD20-Sm147, CD16-Nd148, CD127-Sm149, CD1c-Nd150, CD123-Eu151, CD66b-Sm152, PD1-Eu153, CD86-Sm154, CD27-Gd155, CCR5-Gd16, CD117-Gd158, CD25-Tb159, CD15-Gd160, CD56-Dy161, CD169-Dy162, CRTH2-Dy163, CD371-Dy164, CCR6-Ho165, CD25-Er166, CCR7-Er167, CD3-Er168, CX3CR1-Tm168, CD38-Er170, CD161-Yb171, CD209-Yb172, CXCR3-Yb173, HLADR-Yb174, CCR4-Yb176, CD11b-Bi209.

### Immunohistochemistry

Fibroblasts were seeded in 8-well chamber slides (Ibidi 80826) and stimulated the following day with 1000u/mL IFN-alpha for 30 minutes or 12hrs. The cells were fixed and permeabilized (BD 554714), then stained with DAPI and antibodies against STAT2 (EMD 06-502, 1:200). The samples were evaluated on the Leica DMi8.

### Flow Cytometry

Transduced U6A cells were stimulated for 15-minutes with IFNα2b then immediately fixed with 4% PFA for 10 minutes. Cells were then fixed/permeabilized in 90% ice-cold methanol and stained with anti-phospho-STAT1 (Cell Signaling 9167), a fluorescent secondary antibody (Thermo Fisher) and a LIVE/DEAD viability dye (Thermo Fisher). Flow cytometry was acquired on a BDFACSCanto II and data was analyzed on FlowJo.

### SiMoA Digital ELISA

Plasma samples from the patient and healthy donors were isolated from ficoll-gradient and subsequently clarified by centrifugation at high speeds. Interferon alpha levels were then quantified by digital ELISA using the IFNα Simoa Assay Kit (Quanterix, 100860) according to the manufacturer’s instructions on a Simoa HD1 Analyzer.

