## Supplemental Materials for "Homozygous *STAT2* gain-of-function mutation by loss of USP18 activity in a patient with type I interferonopathy"

### Supplementary Materials

#### Supplementary Table 1: Patients' immunological phenotype from the analysis of whole blood samples.

| | **P1 (**4 months) | Normal range |
|---|---|---|
| CD3+ | 6,700 | 2,500-5,900/$mm^3$ |
| CD3+CD4+ | 3,140 | 1,400-4,300/$mm^3$ |
| CD3+CD8+ | 625 | 500-1,700/$mm^3$ |
| CD16+CD56+ | 190 | 160-950/$mm^3$ |
| CD19+ | 3,310 | 300-3,000/$mm^3$ |

#### Supplementary Table 2. Immunoglobulin levels.

| | **P1 (**4 months) | Normal range |
|---|---|---|
| IgG (g/l) | 11.3 | (7.00-16.00) |
| IgA (g/l) | 0.51 | (0.70-4.00) |
| IgM (g/l) | 1.05 | (0.40-2.30) |

#### Supplementary Table 3. List of variants identified after analysis of the WES of the patient

| **Chromosome** | **Genomic DNA position (GRCh17)** | **ID** | **Reference allele** | **Alternative allele** | **Predicted aminoacid change** | **Gene** |
|---|---|---|---|---|---|---|
| 3 | 75786421 | . | G | A | p.Pro785Ser | *ZNF717* |
| 4 | 165876149 | . | A | G | p.Ser209Pro | *TRIM61* |
| **12** | **56749255** | **.** | **C** | **T** | **p.Arg148Gln** | ***STAT2*** |
| 12 | 57604564 | rs763058234 | C | T | p.Ala4273Val | *LRP1* |
| 12 | 104134424 | rs372110731 | GTC | G | p.Leu1925fs | *STAB2* |
| 12 | 104134430 | rs764116330 | ACAAC | A | p.Asn1927fs | *STAB2* |
| 14 | 105418034 | rs201228544 | G | C | p.Pro1252Ala | *AHNAK2* |
| 17 | 26937539 | . | T | G | p.Asp397Ala | *SGK494* |
| 19 | 761638 | . | A | ACC | p.Ser644fs | *MISP* |

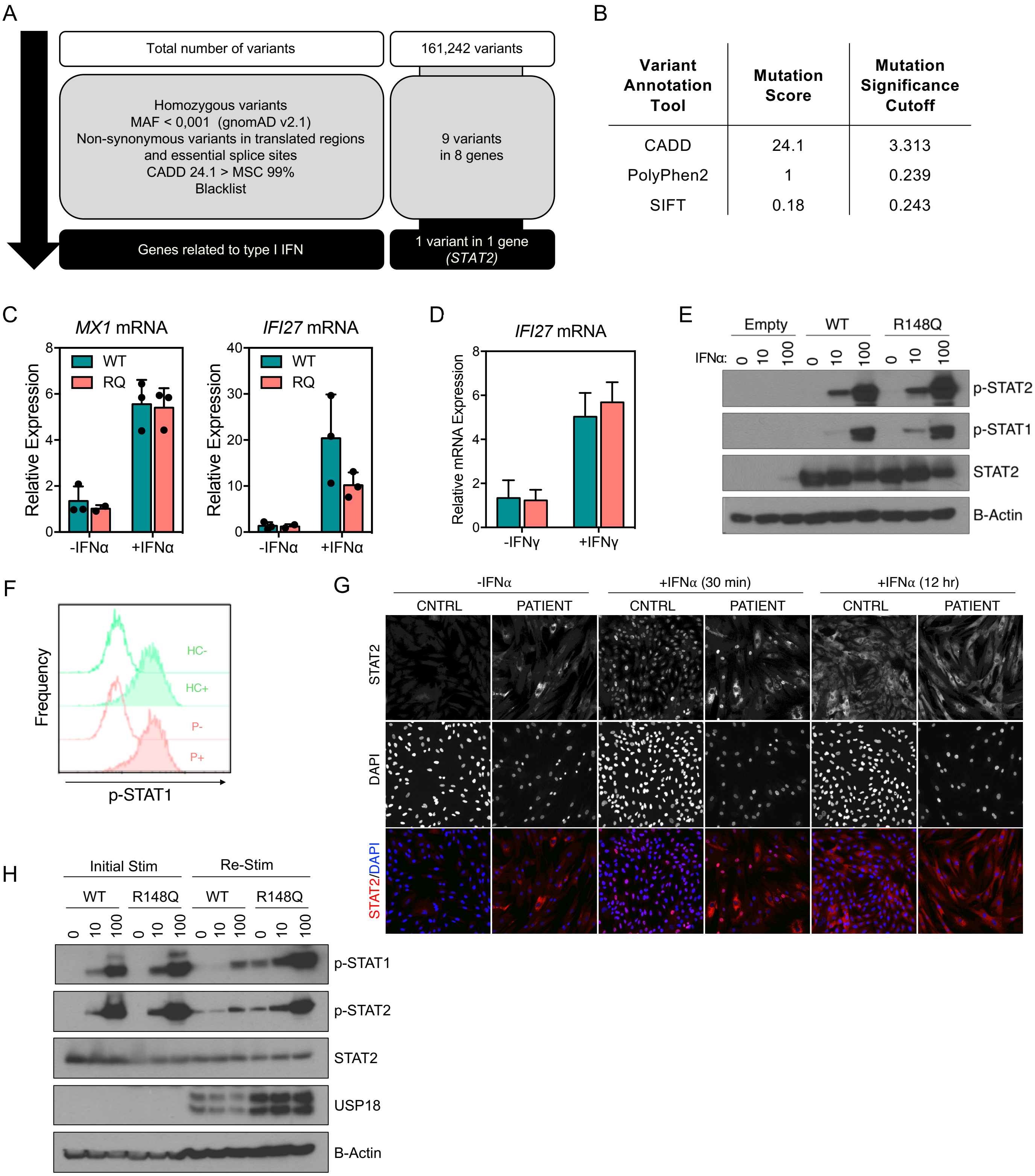


**Supplementary Figure 1**. (A) Schematic demonstrating variant analysis pipeline and results. (B) Prediction of functional impact of R148Q on STAT2 protein. (C) Stimulation of transduced U6A cells with Type I IFN for 4 hours. (D) Induction of target genes by Type II IFN stimulation for 16 hours in transduced U6A cells. (E**)** Analysis of STAT1 and STAT2 phosphorylation in transduced U6A cells following 15-minute stimulation with indicated doses (IU/mL) of IFN-α. (F) Flow cytometry for STAT1 phosphorylation in U6As with WT (HC) or R148Q (P) STAT2, stimulated (+) or not (-) with IFN-alpha for 15 minutes. (F) Immunofluorescence staining of STAT2 translocation in fibroblasts after 30-minute and 12-hour stimulation with 1000 IU/mL Interferon-alpha. (G) Analysis of negative regulatory capability by stimulating cells which had previously been stimulated (re-stim) or not (naive) with a primary stimulus of IFN-alpha for 12 hours, wash, and allowed to rest for 6 hours (rest) or 36 hours before a re-stimulation with indicated doses of IFN-alpha. All results are representative of 2-3 independent experiments.

### Case Report

We investigated one patient from a consanguineous family (second degree) from Morocco (Figure 1A, 2A). Patient (P1, II.9) was a boy born in November 2015 at 38 weeks of gestation, without any problems during the pregnancy. At birth, he received the BCG vaccine, without secondary effects. At the age of 15 days, he presented with diarrhea, oral thrush and axillar and inguinal adenitis. At the age of 22 days, he was hospitalized due the fistulation of adenitis (Figure 2A); cultures of samples were positive for *Klebsiella* extended-spectrum B lactamase (ESBL) enzymes and *Pseudomonas aeruginosa*; direct staining for acid fast bacilli (AFB) was negative (Figure 1A). Histology revealed reactive lymphoid hyperplasia. He received treatment with gentamicin and ceftriaxone but rapidly it was replaced by amikacin, ceftazidime and imipenem resulting in clinical improvement. However, during this hospitalization he had two other findings marked by bronchiolitis and seizures unrelated to the fever (Figure 2A). Cerebral computer tomography (CT) revealed brain bilateral calcifications distributed in the frontal and parietal regions (Figure 2A). Laboratory tests revealed anemia and mild leucocytosis. Immunoglobulins levels of IgG, IgM and IgA were according to the age of patients. Immunophenotyping of T, B and NK cells, as well dihydrorhodamine 123 (DHR) assay on neutrophils, were normal at that time (Supplemental Table 2). He was assumed to have an inborn error of immunity. At the age of 5 months, he was hospitalized for respiratory manifestations of disease. Chest radiographies revealed cardiomegaly, bilateral opacities of the lungs with an interstitial pattern, and absence of pleural effusion. Chest CT scan was suggestive of pulmonary alveolar proteinosis, but bronchoalveolar lavage fluid (BALF) was not performed. PCR for cytomegalovirus was negative. Culture of blood was negative for pyogenic bacteria, mycobacteria and fungi. However, he received multiple antiviral and antifungal drugs, antibiotics including those for mycobacterial infections and steroids. He died during this hospitalization due to the aggravation of pulmonary disease leading to respiratory insufficiency. The family had previously lost two infant siblings, a girl born in 1991 and a boy born in 1992. Both infants had fistulizing adenitis and they died from disease which appeared infectious in origin, but suggestive of uncontrolled inflammation. Other siblings (born in 1994, 1996, 2002, 2005 and 2008 respectively) are healthy and have never been referred for severe infectious diseases or seizures. All individuals received BCG vaccine at the birth according to the Moroccan calendar of vaccines and none had adverse effects.

The patient was referred to the Laboratory of Human Genetics of Infectious Disease, France, for genetic study. Parents signed an informed consent form, in accordance with the requirements of the institutional review boards (IRB) of the various institutions involved. Approval for this study was obtained from the French Ethics Committee (CPP) and the French National Institute of Health and Medical Research (INSERM; 2010-A00650-39); and from the Rockefeller University (JCA-0699).
